# Emergence and maintenance of stable coexistence during a long-term multicellular evolution experiment

**DOI:** 10.1101/2023.01.19.524803

**Authors:** Rozenn M. Pineau, David Demory, Eric Libby, Dung T. Lac, Thomas C. Day, Pablo Bravo, Peter J. Yunker, Joshua S. Weitz, G. Ozan Bozdag, William C. Ratcliff

## Abstract

The evolution of multicellular life spurred evolutionary radiations, fundamentally changing many of Earth’s ecosystems. Yet little is known about how early steps in the evolution of multicellularity transform eco-evolutionary dynamics, e.g., via niche expansion processes that may facilitate coexistence. Using long-term experimental evolution in the snowflake yeast model system, we show that the evolution of multicellularity drove niche partitioning and the adaptive divergence of two distinct, specialized lineages from a single multicellular ancestor. Over 715 daily transfers, snowflake yeast were subject to selection for rapid growth in rich media, followed by selection favoring larger group size. Both small and large cluster-forming lineages evolved from a monomorphic ancestor, coexisting for over ~4,300 generations. These small and large sized snowflake yeast lineages specialized on divergent aspects of a trade-off between growth rate and survival, mirroring predictions from ecological theory. Through modeling and experimentation, we demonstrate that coexistence is maintained by a trade-off between organismal size and competitiveness for dissolved oxygen. Taken together, this work shows how the evolution of a new level of biological individuality can rapidly drive adaptive diversification and the expansion of a nascent multicellular niche, one of the most historically-impactful emergent properties of this evolutionary transition.

## Introduction

Earth’s biosphere looks quite different now than it did a billion years ago, due in no small part to the expansion and success of multicellular organisms^1, 2^. It remains difficult to disentangle whether the evolution of multicellularity itself favors ecological diversification, or whether this is just a consequence of novel multicellular traits evolved by these lineages. For example, the ability for green algae to grow on land, or more appropriately, in the air (the only lineage of the dozens of independently evolved multicellular algae we call ‘plants’), opened up vast new ecological and evolutionary frontiers^3^. Yet this diversification arguably has more to do with the evolution of multicellular traits allowing for exploration of a new environment^4^, including a hydrophobic cuticle, roots, vascular system for transporting water, etc.^5–7^, than it does the transition to multicellularity (the origin of simple multicellular groups capable of adaptation), *per se*^8^.

John Bonner argued that, far before the evolution of groundbreaking multicellular innovations, multicellular size was a trait that favored open-ended ecological diversification^9^. In this view, far before the evolution of trade-off breaking multicellular innovations that characterize many dominant multicellular clades^10^, multicellularity itself offered access to novel size niches, driving ecological divergence and maintaining diversity. Bonner’s hypothesis, while foundational, has remained largely conceptual. This is not uncommon when it comes to theories about the evolution of multicellularity— all well-studied extant lineages made this transition deep in the past, and relatively little is known about the ecological or evolutionary dynamics of their earliest multicellular ancestors.

In this paper, we examine the link between multicellularity and diversity via a combination of long-term experimental evolution and mathematical modeling. We show that, consistent with Bonner’s ‘size-niche’ hypothesis, the simple act of creating a multicellular group generates novel size-dependent trade-offs that drive ecological divergence and maintain coexistence. For this work we used the ‘snowflake yeast’ model system of diffusion-limited multicellularity^11, 12^. Snowflake yeast are an experimental model of undifferentiated multicellularity capable of open-ended laboratory evolution^17^. They have an emergent multicellular life cycle, in which clusters form via incomplete cell division, so that mother and daughter cells remain attached after mitosis, and fracture between cells results in the production of clonal multicellular propagules^13^. The size to which snowflake yeast can grow is an emergent property of the shape of the cells in the group: more elongate cells increase the amount of free space within the cluster interior, increasing the size to which they grow before strain arising from cellular crowding breaks a cell-cell bond, causing the group to fragment^12, 14^. Group size at fracture is highly heritable (broad sense heritability ~0.8, on par with clonally-reproducing animals like *Daphnia*)^15, 16^, and is a trait under strong selection in our system^12, 17^.

In our ongoing Multicellularity Long Term Evolution Experiment (MuLTEE), we use three different treatments (anaerobic, mixotrophic, and obligately aerobic, referred to as PA, PM and PO) to test the effect of metabolic differences on the evolution of multicellularity. To favor the evolution of larger groups, we perform a round of settling selection (using sedimentation speed through liquid media to screen for larger groups), after 24 h of growth in batch culture (Figure 1A). The fitness of snowflake yeast thus depends on their ability to both (i) compete for resources during batch culture, and (ii) form large groups that have high multicellular survival during settling selection. These traits appear to trade-off, as large size reduces the potential for resources to diffuse into the cluster interior and reach internal cells, slowing growth rates, while increasing settling speed and thus survival^11, 12^. Such trade-offs may be common during the early stages of the evolutionary transition to multicellularity. Group size is a key trait under selection in many multicellular lineages, as distinct benefits of multicellularity (*i.e*., protection from predators or harsh environments, motility, or cooperative metabolism^18^) often function in a size-dependent manner. Yet prior to the evolution of circulatory systems or other morphological innovations that mitigate this constraint^19^, larger size will reduce a group’s surface area to volume ratio, potentially causing key exogenous resources (*e.g*., reduced carbon, dissolved oxygen) to become diffusion-limited, slowing growth^18, 20, 21^. In our experiment, glucose is initially present at approximately 4 orders of magnitude higher concentration than oxygen, and poor oxygen diffusion into the cluster interior imposes a strong constraint on the evolution of larger group size^12^.

**Figure 1.**
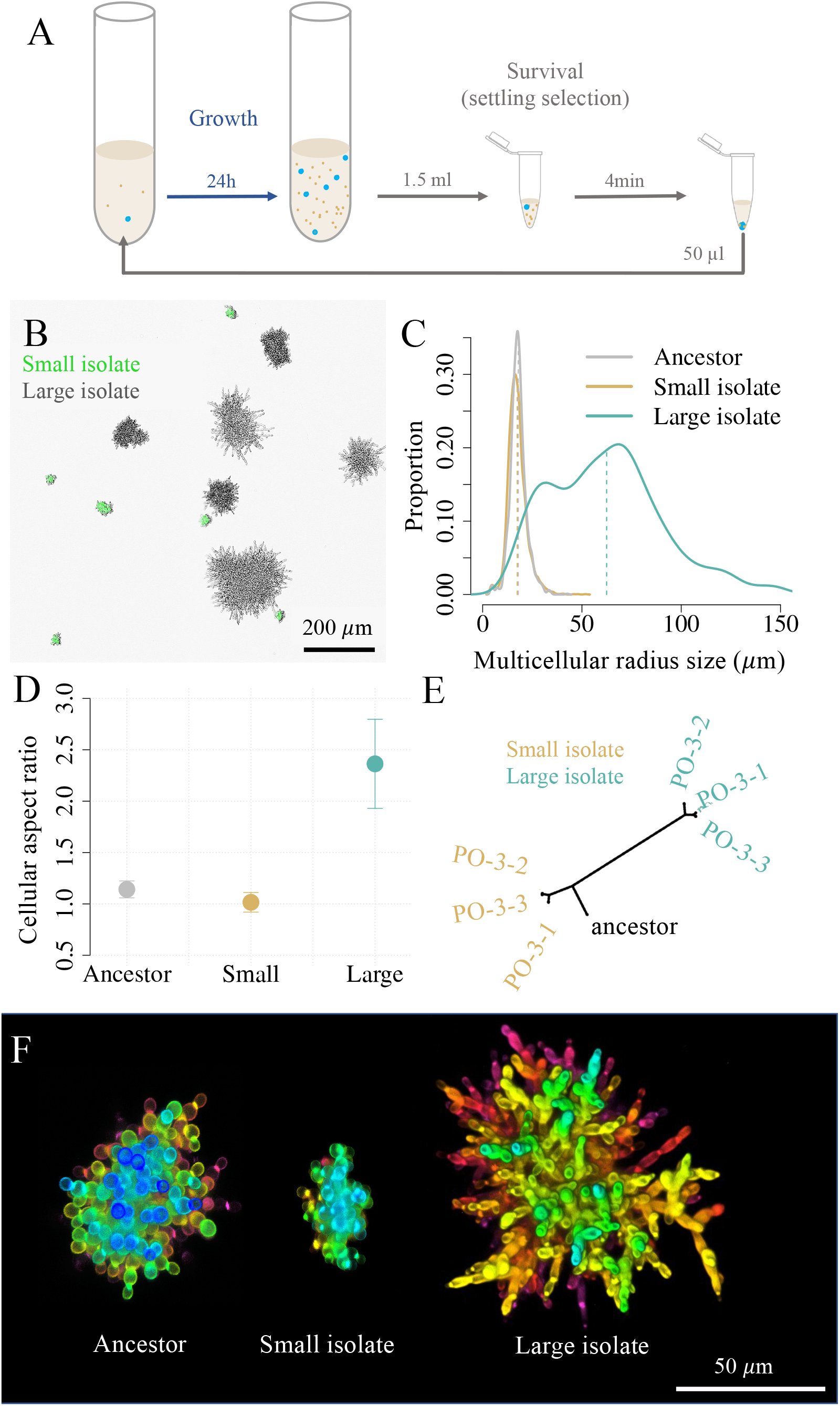
Emergence and long-term coexistence of large and small snowflake yeast phenotypes. (A) Daily transfers consist 24h of batch culture, in which selection favors faster growth, followed by a round of settling selection for larger group size. (B) While the experiment started from a monomorphic multicellular ancestor, after 715 rounds of selection, the population is composed of large and small (GFP for ease of identification) phenotypes. (C) We measured the cluster size distribution via microscopy. Small-sized snowflake yeast (yellow) are similar in size to their ancestor (gray). Large-sized snowflake yeast isolates (teal), in contrast, are 48 times larger. (D) The Large genotype evolved highly elongate cells, with a mean aspect ratio (length to width) of 2.36, while the Small genotype became nearly perfectly spherical (aspect ratio 1.01) from the ancestor’s slightly oblate cells (aspect ratio 1.14). Bars represent one standard deviation. (E) The phylogeny of Small and Large genotypes shows they do not share any mutations, demonstrating that the lineages leading to each have been coexisting for the duration of our long-term evolution experiment. (F) Differences in cellular morphology between the multicellular ancestor, Small and Large genotypes shown via confocal microscopy. Note that the Large cluster shown here is smaller than its maximum possible size (this cluster is in the 40th percentile of size). Color indicates depth.

In prior work within a 60-day evolution experiment^11, 22^, we observed what appeared to be nascent steps in the diversification of snowflake yeast populations into small and large sized strains. Here, we examine what happens when such an experiment is prolonged, and demonstrate that coexistence between small- and large-sized strains is evolutionarily stable, lasting 715 daily transfers (~4,300 generations). We show that size dependent trade-offs involving access to dissolved oxygen are essential for maintaining diversity: coexistence is lost if we provide supplemental oxygen, and never evolved in mixotrophic and anaerobic populations of the MuLTEE capable of fermentation. Coexistence is the result of ecological specialization: small snowflake yeast evolve to become growth-rate specialists, while large group forming genotypes, which are 16 to 48-fold larger, evolve to become survival specialists. Through both experiments and modeling, we show that coexistence is maintained by frequency-dependent selection, and that the large difference in group size between size specialists is the result of niche partitioning. Taken together, this work demonstrates that a simple and yet fundamental trade-off between growth and survival, mediated by differential oxygen diffusion through bodies of different sizes, can drive and maintain ecological diversity in a nascent multicellular lineage.

## Results

### Maintenance of two distinct phenotypes over long-term evolution

After 715 serial passages with daily selection favoring faster growth and larger group size (Figure 1A), three of five populations in the obligately aerobic treatment (PO) appeared to contain a mixture of small and large snowflake yeast (PO-3, PO-4 and PO-5, Supplementary Figure 1; see Supplementary Figure 2 for microscopy images of population PO-4 after 715 serial passages). To determine if these size differences reflect heritable variation among coexisting strains, or reflect within-genotype phenotypic variation (*i.e*., small groups may simply be fragments of large group-forming genotypes), we isolated 10 small and 10 large snowflakes from each of the five replicate populations with a dissecting microscope, and then measured the size of the resulting populations after growth. In the three populations showing clear size heterogeneity (PO-3,4,5), the populations derived from large clusters were 37, 27, and 16-fold larger than those derived from small genotypes, respectively, while they were only 2 and 1.5-fold larger in the two populations lacking visible signs of heterogeneity (populations PO 1&2, Figure 1B; Supplementary Figure 1). This was surprising, both because our detailed investigations of the anaerobic (PA) and mixotrophic (PM) treatments of the MuLTEE did not show any sign of size dimorphism^17^, and because daily size selection should, in theory, favor the larger, faster-settling genotypes.

To understand the drivers of coexistence between small and large-sized snowflake yeast, we isolated representative genotypes from the population where we first noticed the size dimorphism (PO-4) after 715 transfers (Figure 1B and F). We chose this time point because the MuLTEE was on transfer 715 when we began investigating this phenomenon, and we wanted to use the most highly-evolved populations available. Clusters of the small genotype (henceforth called Small) were similar in size to the ancestor of the experiment, with a mean radius of 17.4 μm (vs. 17.9 μm for the ancestor, *n* = 1454 for ancestor clusters, *n* = 2860 for small clusters, Tukey’s HSD after one-way ANOVA, F_2,5085_ = 11255, *p* = 0.1 for the post-hoc test). In contrast, the larger genotype (henceforth called Large) evolved to form clusters with a mean radius of 62 μm, a ~48-fold increase in volume from the ancestor (Figure 1C, *n* = 774 for large clusters, *p* < 0.001, Tukey’s HSD for the above ANOVA). Prior work has shown that snowflake yeast primarily evolve larger size by increasing the length of individual cells (increasing their aspect ratio), which in turn reduces cellular packing density and allows clusters to grow larger before fracturing^14, 17^. We examined whether evolved differences in aspect ratio were responsible for differences in the Small and Large cluster phenotypes. Consistent with prior experiments, cells within Large clusters evolved to be more than twice as elongated as the ancestor, increasing their aspect ratio from 1.1 to 2.3 (Figure 1D and F, *n* = 128 for ancestor cells, *n* = 120 for large cells, Tukey’s HSD after one-way ANOVA, F_2, 295_ = 95.5, *p* < 0.0001). In contrast, cells in the Small snowflake yeast genotype evolved to be nearly spherical, with their mean aspect ratio declining from 1.14 to 1.01 (*n* = 50 for small cells, *p* = 0.02, Tukey’s HSD for the above ANOVA).

To determine if the evolution of Small and Large phenotypes is recent, or occurred early in the experiment, we sequenced the genome of three randomly-selected isolates of each type, from population PO-4 after 715 transfers. We found fifteen mutations shared among the Small isolates, four of them linked to respiration (Supplementary Table 1). Sixteen mutations were shared among the Large isolates, four of them are involved in the cell cycle, growth, or cell wall organization, and are thus candidate mutations for increasing cellular aspect ratio^12, 17^ (Supplementary Table 1). However, no mutations were shared between Small and Large genotypes (Figure 1E and Supplementary Table 1). Because this experiment was started by a single clone, this indicates that the last common ancestor of the two phenotypes is the monomorphic ancestor of the MuLTEE. The lack of shared *de novo* mutations further suggests that lineages leading to these distinct phenotypes diverged early on in the MuLTEE and have been coexisting for ~4,300 generations^17^ of growth and 715 rounds of size-based settling selection.

### Competition for oxygen creates a niche for coexistence

Frequency-dependent selection, in which the fitness of a genotype declines/increases as it becomes relatively more common/rare in the population, can maintain coexistence even in the face of stochastic perturbations to equilibrium genotype frequencies^23, 24^. To determine if frequency-dependent selection is acting to maintain a polymorphic population of Small and Large genotype snowflake yeast, we competed these strains across a wide range of starting frequencies, from 1% to 80% Large (Figure 2A). In these experiments we used the same conditions as in the MuLTEE: a growth phase of 24 hours in YPG media, followed by a survival phase where we select for larger size (Figure 1A). Regardless of their starting frequency, all populations evolved to have an average of 9% Small snowflake yeast after 6 rounds of growth and selection. This frequency was nearly identical to that of the population as a whole from the t715 PO-4, where large genotypes of yeast were present at a mean of 9.4% (Supplementary Figure 2). The fitnesses of both Small and Large snowflake yeast are strongly frequency dependent. We estimate that the two strains possess equal fitness values when the Large strain composes 9% of the population and the Small strain composes the other 91% (Figure 2B).

**Figure 2:**
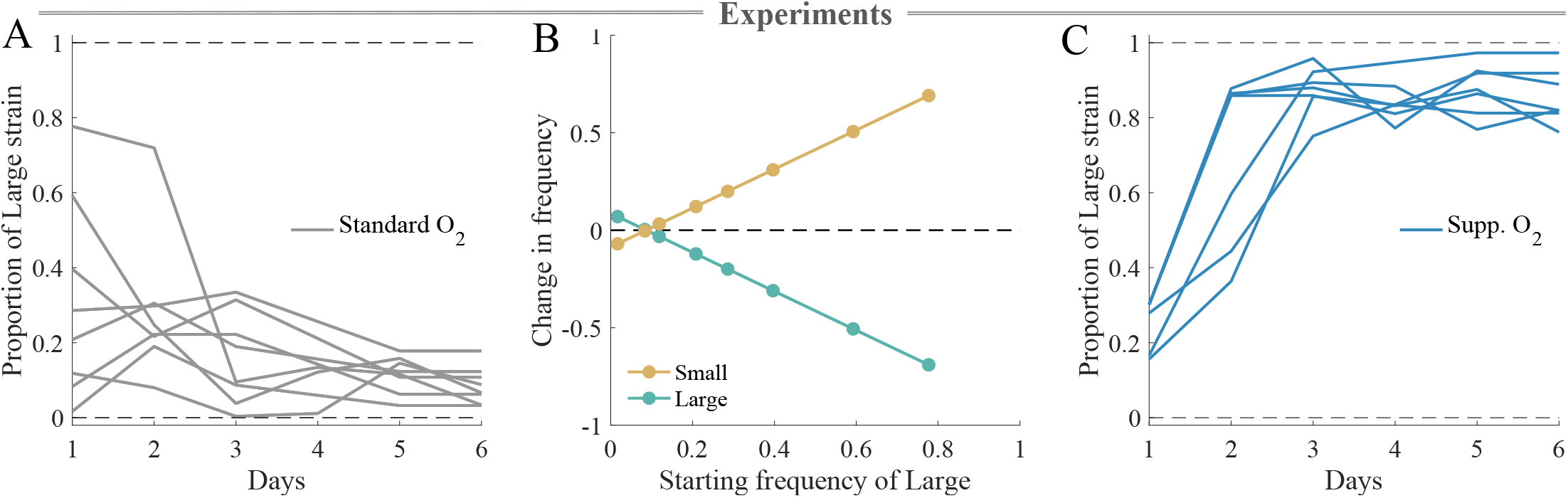
Coexistence between Small and Large group forming genotypes is mediated by oxygen. (A) Under standard oxygen conditions (~26% PAL, gray) our populations converge on an equilibrium of ~9% ± 4.8. Large snowflake yeast regardless of their starting frequency (*n*=8). (B) Under standard oxygen conditions, frequency-dependent selection maintains a co existence between Small and Large strains at 9% Large clusters. Data points reflect changes in frequency of each genotype over 6 days of culture from the experiment shown in A; fitted lines are simple linear regressions. (C) The Large genotype has an advantage under supplemental oxygen, increasing to an equilibrium frequency of 86% ± 7.3 in a high-oxygen environment (~84% PAL, dark blue, *n*=6).

Snowflake yeast in this experimental treatment are obligate aerobes and do not possess the ability to actively transport oxygen throughout their tissues. Instead, oxygen diffuses passively through the cells and the media surrounding them, and diffusion limitation is a fundamental constraint on the growth of interior cells^12^. Prior work has shown that in this system, respiration consumes oxygen faster than it can diffuse into the media, resulting in a mean partial pressure of oxygen at ~26% of present atmospheric levels (26% PAL) under our standard growth conditions^12^. We thus hypothesized that, due to their smaller size and reduced diffusion limitation, Small clusters may have a competitive advantage for O_2_, allowing them to be maintained at a high equilibrium frequency despite daily settling selection favoring large size. To determine whether the coexistence equilibrium is sensitive to oxygen, we performed a competition with supplemental oxygen, provided via an in-tube aerator (which prior work has shown increases mean *p*O_2_ from ~26% to ~84% PAL in the growth medium^12^). We initiated five replicate populations with intermediate frequencies of the Large genotype (~25%), then passaged them for 5 rounds of growth and settling selection. Rather than declining to the 10% equilibrium as seen in the standard oxygen conditions (Figure 2A), under supplemental oxygen the Large genotype rose to 86% frequency (Figure 2C), indicating that oxygen availability is a key driver of the coexistence equilibrium, with more oxygen favoring a higher frequency of the Large genotype.

### Functional specialization mediated by competition for oxygen and selection for large size

We next investigated the mechanism underlying coexistence via frequency dependent selection. Small and Large snowflake yeast may coexist due to emergent functional specialization over a classic trade-off between key life history parameters: Small snowflake yeast appear to be growth specialists, outcompeting Large genotypes for oxygen, though at the expense of low survival during settling selection. In contrast, Large snowflake yeast may be following a strategy of slow growth but high survival. To examine this hypothesis, we developed a mathematical model with two steps: a growth phase, and a survival phase (Figure 3A). The Small (S) and Large (L) clusters grow given a maximum rate *μ* modulated by resource availability in a two-resource Monod growth equation^25^, and die at a rate *δ*, representing the daily rate of background cellular mortality of cells in snowflake yeast^13^ (Figure 3A, see Methods for the equations). As obligate aerobes growing through most of the culture cycle with an abundance of sugar, the growth rate of yeast cells depends on the limiting resource, oxygen (R), which is both present at the start of the growth phase and can be added at various rates through its duration (Figure 3A).

**Figure 3:**
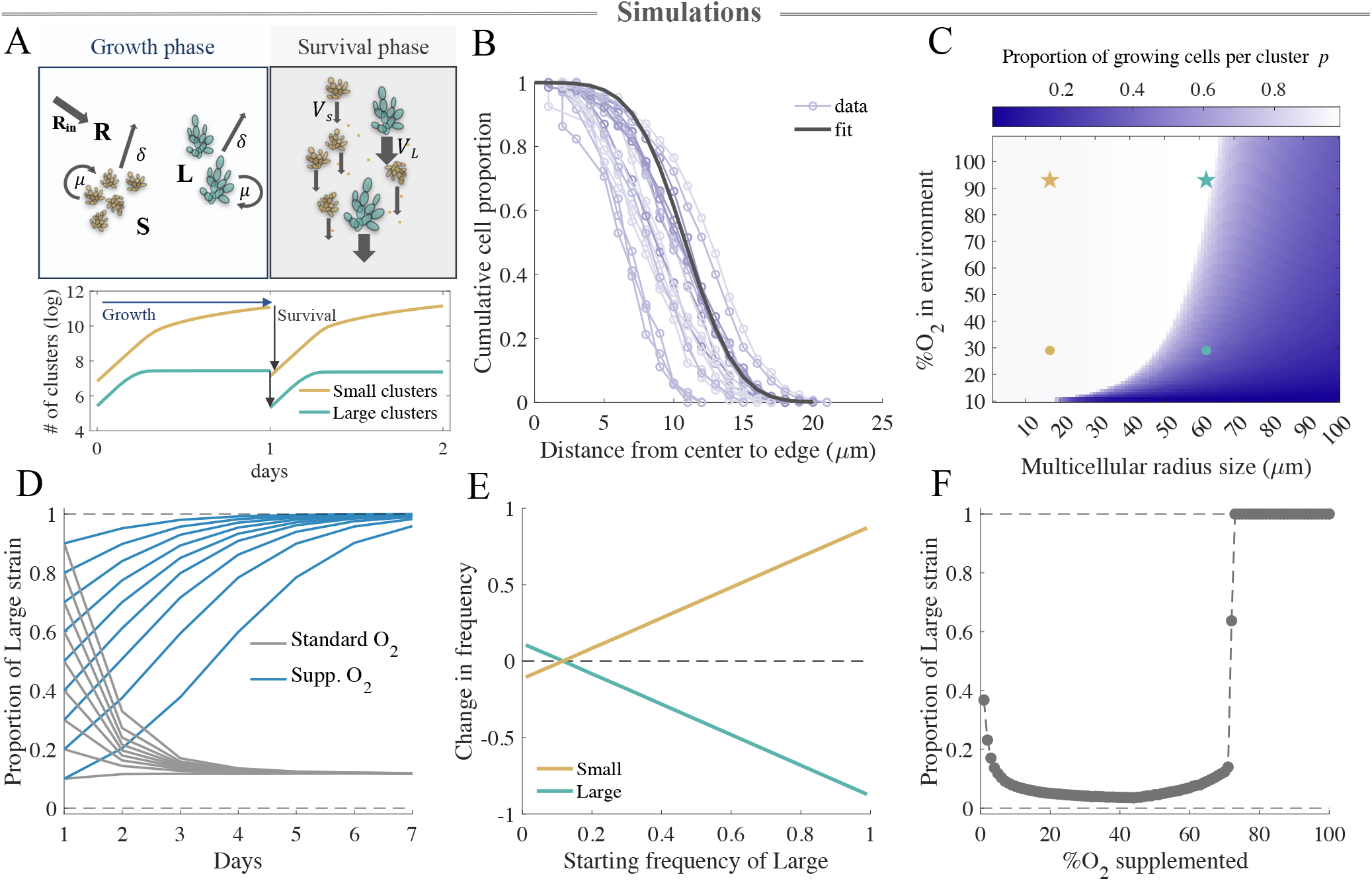
Modeling coexistence. (A) To examine the dynamics of competition between Small and Large snowflake yeast, we developed a model with two phases: growth, in which Small (S) and Large (L) clusters grow by respiring oxygen (R), and survival, that starts at the end of the growth phase and selects a proportion of the total population based on their settling speed (*V)*. The bottom panel shows the growth dynamics of each cluster type followed by settling selection. (B) We use a normal distribution fit to approximate the distribution of cells within individual snowflake yeast based on scanning electron microscopy data (SBF-SEM) (Supplementary Figure 4). (C) By utilizing the distribution in (B), we calculate the proportion of cells in the cluster that have access to oxygen and can grow, across a range of cluster sizes and oxygen concentrations. The yellow and teal markers correspond to the Small cluster size and the Large cluster size for the mean oxygen concentrations under standard (dots) and high oxygen conditions (stars). (D) The model recapitulates the experimental dynamics (Figure 2), with standard oxygen conditions supporting an equilibrium of 11% of Large snowflake yeast, and supplemental oxygen favoring the Large strain. E) Much like the experiment (Figure 2C), coexistence in the model is mediated by frequency-dependent selection, with a stable equilibrium at 11% Large. (F) We investigated the role of oxygen availability on the outcome of competition in the model by supplementing our populations with oxygen (added hourly, and capped at 100% *p*O_2_). Coexistence in the model is dependent on relatively low oxygen availability and is lost if the concentration of supplemented oxygen exceeds 73%.

Oxygen gets depleted as the populations grow, and its concentration sets the distance within the cluster that oxygen can diffuse and be used by yeast cells for growth. To estimate the proportion of the cluster that has access to oxygen and can grow, we model the diffusion of oxygen through the cluster. We find from high-resolution serial bulk face scanning electron microscopy that cells are more densely packed near the core of the cluster, and more loosely packed towards the edge. We model this distribution of cell positions using a normal distribution (cumulative distribution function, Figure 3B). We use this estimation to model the heterogeneous diffusion of oxygen in clusters for a wide range of sizes and oxygen concentrations, from which we deduce the proportion of cells in a given cluster that have sufficient oxygen to grow (termed *p* in the growth equation, see Methods, Figure 3C).

At the end of the 24-hour growth phase, we simulate settling selection. We randomly position a subset of our population within the tube, then use Stokes’ law^26^ to calculate the size-dependent terminal settling velocity of each cluster (Figure 3A). As in the experiment, settling selection favors larger clusters, but stochasticity in starting position (some clusters start out near the bottom regardless of their size) allows some small groups to survive. In this model, the only difference between the two competitors is their size (we used radii of 17 μm and 62 μm for Small and Large, respectively, based on their average size in our experiments, see Figure 1C). Hence, fitness depends just on size-dependent growth, which is mediated by oxygen diffusion during the culture phase, and size-dependent survival during the settling selection phase.

Our model recapitulates key dynamics observed in the experiment: competition over oxygen, the primary resource determining growth rate, maintains coexistence with a ~11% frequency of Small (Figure 3D&F) through frequency-dependent selection (Figure 3D). Confirming the critical role of oxygen limitation for maintaining coexistence, Large competitively excludes small at sufficiently high O_2_ concentrations, achieved by oxygen supplementation (Figure 3F and Supplementary Figure 3). This differs from our experiment (Figure 2C), where adding supplemental oxygen increased the equilibrium frequency of Large dramatically (from 10% to 86%), but not to the point where the Small genotype was entirely displaced.

Both our experimental competition and model show that the fitness of Small and Large snowflake yeast is frequency dependent, leading to the stable coexistence of both strains (Figure 2C and 3E). We used our model to examine how selection acting on size dependent growth rate and survival depends on genotype frequency (Figure 4). Specifically, we varied each genotype’s initial frequency and examined their fitness during the growth and settling phases of the culture cycle. Small has a clear growth advantage, which diminishes as it becomes more common (Figure 4A). This makes sense: smaller groups grow faster than large groups, due to having a larger proportion of their cells oxygenated. When the population is dominated by the slower-growing Large genotype, it takes longer for them to reach stationary phase. The less frequent Small is in the population, the more hours it has to compound its growth advantage, gaining a frequency dependent growth advantage.

**Figure 4:**
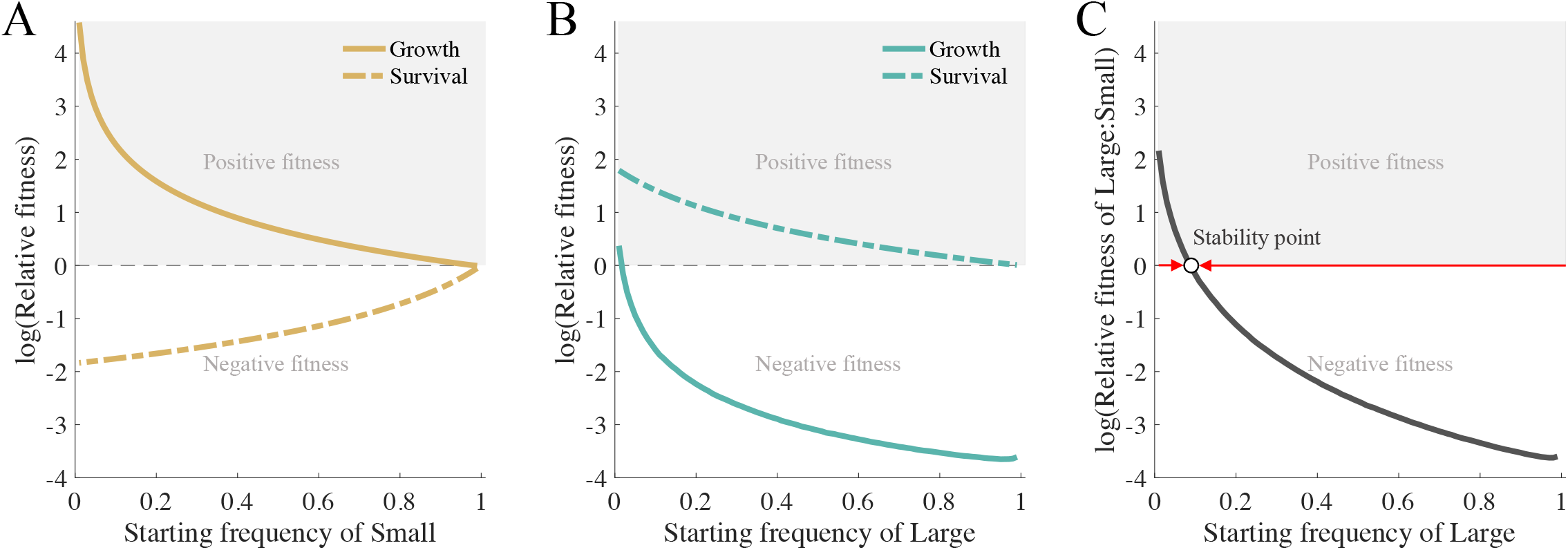
Frequency dependence and specialization along a growth-survival trade-off. We calculate the relative fitness of Small and Large clusters between the start and the end of the growth and the survival phases, for a range of initial frequencies. For each genotype, we express fitness as the change in frequency of the focal genotype over a single round of growth or settling selection, and the total fitness across a single culture cycle as the product of these multiplicative factors. (A) Small snowflake yeast are growth specialists, but their advantage declines with their frequency. (B) In contrast, Large snowflake yeast have an advantage during settling selection, but this advantage also declines with increasing frequency. (C) The product of relative fitness during growth and settling selection, relative fitness across the culture cycle, has one stable equilibrium point (white dot), maintaining coexistence.

The situation is inverted for Large. This genotype is specialized in higher group survival, though this too diminishes as it becomes more frequent in the population (Figure 4B). This appears to be due to the nature of settling selection: competition to reach the ‘survival zone’ is fierce, as only a small proportion (1/27th) of biomass is transferred after settling selection. When rare, Large groups primarily displace Smalls, but as they increase in frequency, so does competition among Large groups for survival, making them most competitive when rare. Note that when Large is rare, that is also the situation where Small has the weakest growth advantage. As a result, each strain has its strongest advantage when rare. When overall fitness is computed, there is a single equilibrium point at which both strains coexist (white dot in Figure 4C). This equilibrium is stable, as frequency perturbations in either direction —which do not result in one strain’s extinction— will return to the stability point (Figure 4C).

### Size-dependent selection generates and maintains diversity

To determine the conditions under which our model predicts coexistence between genotypes of different sizes, we simulated all pairwise competitions between strains forming groups with radius 10-100 μm in standard oxygen conditions (see Methods for more details on the algorithm). For each simulation we assumed there is a resident population of a certain size, and we examined the outcome of invasion by another strain with a different characteristic size. There are three possible outcomes in such competitions: 1) the larger genotype displaces the smaller, 2) the smaller displaces the larger, or 3)they reach an equilibrium where both strains coexist (Figure 5). Under our model parameters, coexistence is possible over a large fraction of the state space. We have plotted the size of the small and large genotypes that evolved independently in replicate populations PO-3, PO-4, and PO-5 (Figure 5, stars), which all lie within a similar region of the plot supporting coexistence.

**Figure 5:**
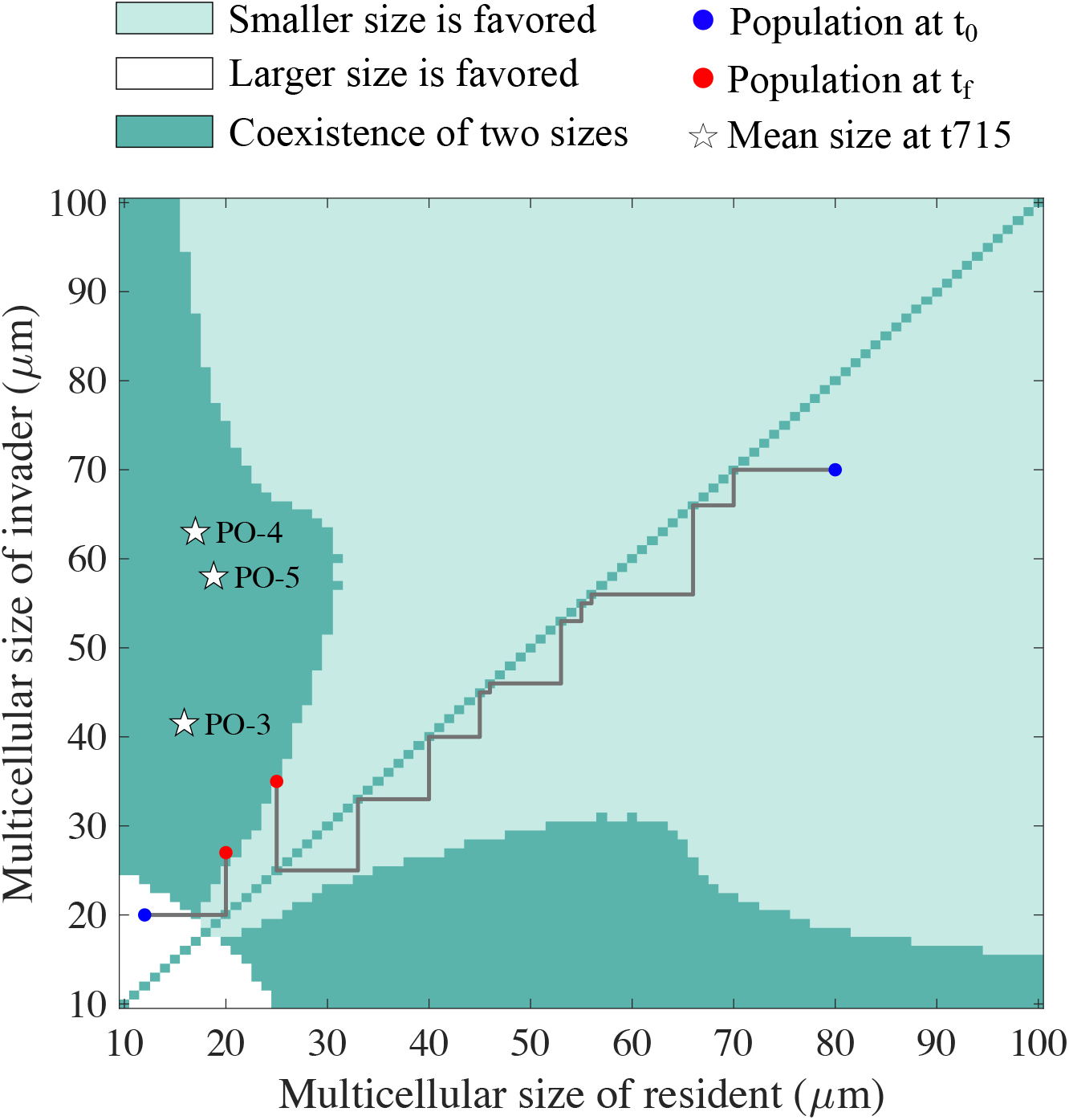
Coexistence is evolutionarily stable. To survey the landscape of ecological interactions between small and large snowflake yeast, we performed an invasion analysis (invader at 1% frequency) of all pairwise comparisons between strains with radii of 10-100 μm. There were three possible outcomes: selection favors coexistence (dark green), the smaller strain wins (light green) or larger strain wins (white). Over this image, we have plotted the trajectories of two populations in which we simulate iterative rounds of mutation and selection (blue and red circles denote start and end points, respectively). This simulation shows how selection can favor directional evolution (either increasing or decreasing size depending on their starting point) until ultimately reaching a portion of the landscape favoring coexistence between small and large sized strains (dark green zone). The white stars correspond to the mean sizes of the Small and Large isolates from the three replicate populations that independently evolved size dimorphism (populations PO-3, PO-4, PO-5) after 715 transfers.

We then explored the trajectory of evolution along this landscape by simulating a series of pairwise competitions, between populations starting out either smaller or larger than those shown to coexist (starting from 12 μm or 90 μm on Figure 5, see other example trajectories in Supplementary Figure 5). In this simulation, novel mutants were generated by drawing from a uniform distribution ± 12 μm centered on the resident strain’s size. Starting the mutant at 1% frequency, we simulated competition until an equilibrium was reached. If either strain displaced their competitor, they became the new resident. In all simulations, selection on this landscape drives snowflake yeast strains that are too large or too small into intermediate sizes that were capable of coexisting through frequency-dependent selection (of the kind described in Figure 2C, Figure 3E, and Figure 4). In our simulations, coexistence is an evolutionarily stable equilibrium, in which the system, if perturbed via the fixation of either a very small or large genotype, will return to a state of coexistence given mutationally-generated novelty.

Interestingly, the Large and Small strains from PO-3, PO-4 and PO-5 are more divergent in size than would be expected via the model (see Figure 5). One possibility is that character displacement has driven further ecological specialization. We tested this hypothesis via experimental evolution (Figure 6). We evolved five replicate populations of the Large and Small genotypes in the absence of an opposite-sized competitor for 40 rounds of growth and settling selection. Each genotype evolved towards an intermediate size in the absence of competitors—with the Small strain evolving to be 1.7 times as large (from 14 ± 1 to 25 ± 3 μm on average, regression slope *β* = 0.27, Figure 6A), while the Large strain evolved to be an average of 15% smaller (from 55 ± 3 to 47 ± 1 μm on average, regression slope *β*=−0.27). The regression slopes were significantly different (the interaction between genotype and time on cluster size, assessed via ANCOVA, *F_(1,38)_*=14.9, *p* < 0.001). We find similar dynamics in our simulations (Supplementary Figure 5). Remarkably, the Small populations even re-evolved a bimorphic population of small and large genotypes in as little as 15 transfers (Figure 6B).

**Figure 6:**
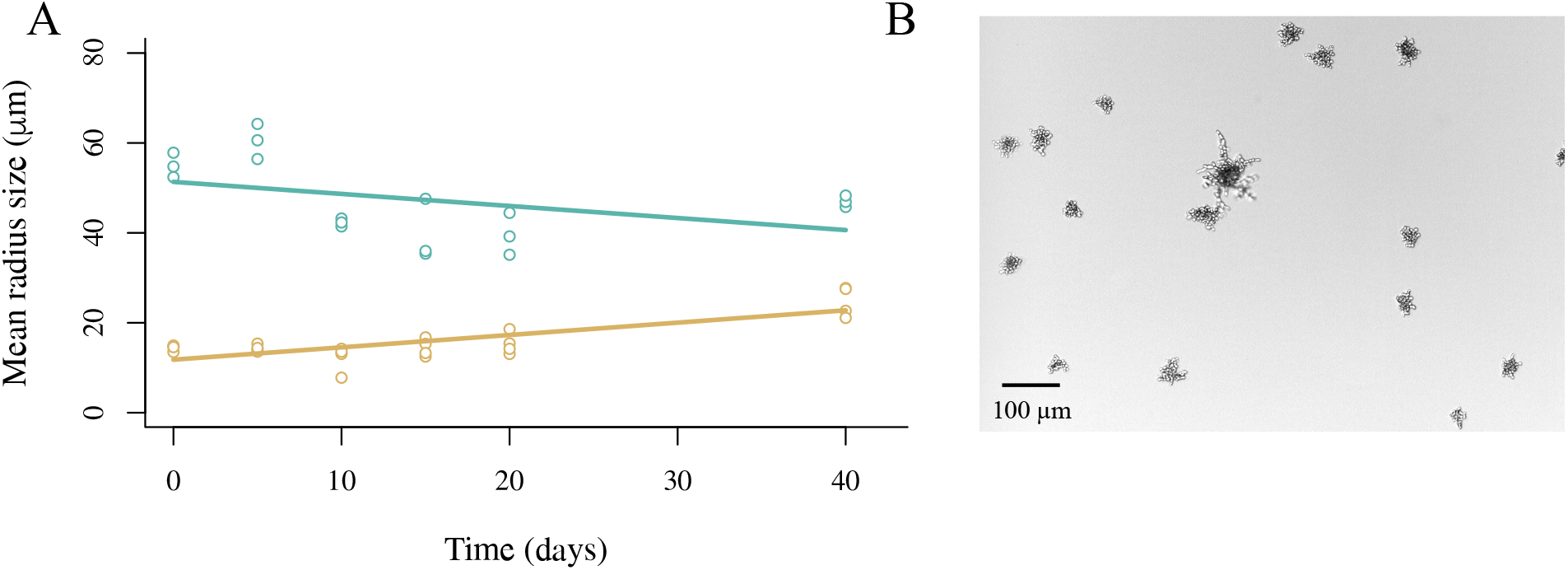
The divergent traits of Large and Small genotypes appear to have arisen via character displacement. (A) To determine if competition between Large and Small snowflake yeast strains have driven divergence due to character displacement, we evolved five replicate populations of the Small and the Large isolates separately for 40 days. Removing their competitor resulted in the rapid evolution of intermediate sizes, with the Small strain evolving to be approximately twice as large (14 μm to 27 μm) while the Large strain shrank on average by 15% (from 55 μm to 47 μm). This suggests the divergence seen between these strains was the result of competitive interactions, with character displacement evolving to minimize competitive overlap at intermediate phenotypes. (B) We observe the re-emergence of small and large strain diversity in the Small monoculture re-evolution experiments in as little as 15 transfers. Shown is an image from a replicate population after 40 days.

## Discussion

The evolution of multicellular organisms from unicellular ancestors precipitated some of the largest and most consequential adaptive radiations in the history of life on Earth^27–31^. Earth’s ecosystems would be fundamentally different without plants, animals, fungi, or seaweeds—a result of the truly biosphere-altering impact of multicellular taxa^32–38^. However, understanding the ecology of contemporary multicellular organisms, in which extensive cellular differentiation enables functional specialization, is fundamentally different from understanding the ecological implications of the first steps in this major evolutionary transition. In this paper, we explore initial steps in this transition, the formation of simple replicating multicellular groups which enable a diffusion-mediated trade-off between growth and survival that drives ecological divergence and maintains coexistence.

Our biphasic selective environment is highly heterogenous, favoring rapid growth during the 24 h culture cycle, followed by strong selection on multicellular size. Theory and prior experiments across diverse systems suggest that temporally varying environments typically favor generalism, because generalists display a higher overall mean fitness across a wider niche than specialists^39^. In our case, the generalist strategy would be an intermediate size that optimizes fitness along the trade-off between growth and survival. Why might we see specialization and coexistence? In our system, there is no genetic constraint on the evolution of a generalist snowflake yeast. Across the MuLTEE, we see genotypes evolving with radii ranging from 17 μm (Smalls in this experiment) to more than 500 μm (macroscopic snowflake yeast in the t600 anaerobic populations)^12^. Niche partitioning could, in principle, drive divergent selection if the population contains different genotypes with even modest, initially stochastic variation in size. Specifically, if the presence of a slightly larger competitor distorts the fitness landscape, such that there is a local fitness minimum just below their size, then smaller strains may adapt along the growth-survival trade-off by evolving to become smaller. One could imagine the same process occurring for genotypes that are slightly above the mean size evolving to become larger survival specialists. Further work will be required to determine if each size specialist competes primarily with the opposite type, or whether diverse populations exist within each size class and competition is mainly within other conspecific specialists. While the convergent evolution of large and small genotypes in 3 of 5 populations suggests that divergence is relatively robust, further work is required to understand the lack of diversity in PO-1 and PO-2. Intriguingly, these two populations are composed of intermediate sized snowflake yeast (Supplementary Figure 1), suggesting a generalist strategy can be feasible.

It is noteworthy that coexistence in our system is dependent on a trade-off between growth and survival fitness. These are fundamental life history traits that arise from the fact that there are only two ways for a reproducing entity to increase its Darwinian fitness: it can increase its frequency in a population by reproducing more or dying less^40^. Trade-offs between increased reproduction and increased survival are nearly universal in biology^41^, as trade-off free adaptations would be expected to have fixed long ago. Adaptation along these trade-offs is well-known to drive the emergence of novel ecological strategies^42, 43^, and it is interesting that selection for such diversification appeared to occur in our system as soon as multicellularity evolved (inferred from the deep coexistence of both Small and Large lineages).

Experimental evolution has shown, in diverse systems, how simple and stable ecological communities can arise from monomorphic origins through a process of niche construction. Similar to our system, oxygen gradients in test tubes of nutrient broth favor the diversification of an initially clonal population of *Pseudomonas fluorescens*, into a ‘wrinkly spreader’ which grows as a biofilm along the surface, and a ‘smooth’ genotype that occupies the low-O_2_ broth below^44, 45^. *E. coli* evolving on glucose minimal media diversify into genotypes that either specialize on this resource, or on overflow metabolites that result from glucose consumption, like acetate^46^. Yeast evolving in static culture tubes differentiate into wall-adherent and bottom-dwelling genotypes, reducing competition caused by cellular crowding^47^. Even RNA replicators evolve into stable ecological networks of hosts and parasites^48^.

Little is known about the joint dynamics of early multicellular adaptation and ecological assembly—though clearly organismal biology is tightly coupled to ecological opportunities^49, 50^. Our results here do not explain the remarkable diversity of modern multicellular ecosystems^51, 52^. Instead, our results focus in on the very first steps in the evolutionary transition to multicellularity—the evolution of replicating groups of cells that are Darwinian Individuals, capable of multicellular adaptation^53^. Our results support a central component of Bonner’s foundational work on the evolution of multicellularity^9, 54, 55^: group size is itself a key trait underpinning the origin of novel niches. Our paper is not the first to show that group formation itself creates a niche-multicellularity has evolved repeatedly in diverse model systems when group formation is adaptive^18^. Here, we build on prior work and use experimental evolution to demonstrate how an emergent trade-off stemming from group size can expand multicellular niche space, from a single niche to two, occupied by growth and survival specialists, potentially resulting in the long-term coexistence of distinct lineages.

This work raises a number of questions of broad interest in the field, most prominently: how does ecological specialization interact with organismal specialization? In the context of MulTEE, we intend to explore mechanisms by which divergent selection on Small and Large lineages results in different forms of morphological or cellular specialization. Moreover, once coexistence between different size-based strategies emerges, it is unclear whether such coexistence is stable indefinitely or whether further multicellular adaptation opens up additional niche space. In theory, the emergence of diverse lineages may enable further niche construction, perhaps through the evolution of novel ecological interactions (as has been found in other evolution experiments^56, 57^), or the collapse of diversity through the evolution of trade-off breaking mutations^58–60^. Exploring these questions will shed light on the process and limits to open-ended diversification once the evolutionary transition to multicellularity enables individuals in complex populations to grow ever larger in size.

## Methods

### Model system

Snowflake yeast are a simple model system of undifferentiated multicellularity, which original evolved due to selection for faster sedimentation^11^ in the diploid unicellular yeast strain Y55. This phenotype was associated with a single mutation in the *ACE2* gene, responsible for incomplete cell separation^16^. For the Multicellular Long Term Evolution Experiment (MuLTEE)^11, 26^, we wanted to study the process of multicellular adaptation without confounding this by having different mutations driving group formation, so we deleted the *ACE2* gene in our ancestor (*ace2*Δ*∷KANMX// ace2*Δ*∷KANMX*). This initial snowflake genotype was used as the basis of three treatments (with 5 independent reps per treatment): obligate anaerobic metabolism (growth on glycerol, which is non-fermentable), obligately aerobic metabolism (via the selection of a spontaneous petite mutant that cannot respire), and mixotrophy, in which yeast both ferment and respire. In this paper, we focus on the obligately aerobic treatment, as that is where we saw coexistence.

### Estimating the number of generations during experimental evolution

In order to estimate the number of generations elapsed over 715 transfers, we used data from Bozdag et al (2021)^17^, in which we measured the biomass in the population by imaging at the beginning and at the end of a 24-hour growth cycle for each of the five independently evolving lines in the MuLTEE at three time points: 200, 400 and 600 days. Next, we calculated the number of generations by taking the log_2_ of each day’s fold increase in biomass, assuming that days 0-200 were described by the t200 generation time measurement, 200-400 was described by the t400 generation measurement, and so on. Using this approach, we estimate that the PO populations underwent ~4,300 generations of growth over the 715 transfers.

### Detecting size polymorphism in t715 evolving populations

At 715 transfers, three out of five replicate populations of obligately aerobic snowflake yeast appeared to be composed of both small and large genotypes. To determine if the small groups present were simply fragments of large group-forming genotypes, or represent genotypes that are unable to form large groups, we isolated 10 small and 10 large clusters from each population using a Singer SporePlay tetrad dissecting microscope. We then grew each of these genotypes in isolation, and measured their size distributions (Supplementary Figure 1).

### Isolation of small and large cluster types

In order to examine the dynamics of small and large strain competition, we isolated a single pair of isolates from line PO-4, which we term Small and Large, respectively. We marked the Small strain with yeGFP (plasmid pFA6a-TEF2Pr-eGFP-ADH1-Primer-NATMX4 amplified and inserted intro *LYS2* locus) using the LiAc/SS-DNA/PEG method of transformation^61^, allowing us to distinguish it from the unlabeled Large genotype during competition experiments. To measure the fitness cost of GFP expression, we labeled the unicellular ancestor (strain GOB8) with the same construct, and competed it against its unlabeled counterpart. The cost was negligible (5 replicates, 2 transfers, t-test *t*=−1.26, *p*=0.25, see Supplementary Figure 6). Representative clusters were imaged using a Nikon AR1 confocal microscope.

### Competition assays

The monocultures for small and large clusters were grown overnight in a shaking incubator (30°C, 250 rpm) in fresh liquid media (YEPG, yeast extract peptone glycerol). The density of each population was estimated before starting the competition with the following procedure: we diluted the overnight cultures 1000-fold and placed 500 μL of the diluted cultures in a 24-well plate. We estimated densities by imaging the populations using Nikon Eclipse Ti inverted microscope, counting cluster number via a custom script in ImageJ. We calculated the volume of starting culture based on the cluster density and the desired starting frequency. The two phenotypes were placed together in fresh media and grown for 24 hours. To calculate the frequency of both genotypes at the start of the experiment, we again used microscopy, differentiating between strains by the presence of a GFP marker (the small genotype was labeled). We performed settling selection from Day 1 cultures and measured the population densities before and after selection to disentangle the growth from the survival advantages of each type. To select for larger size, we placed 1.5 mL of the overnight culture in a 2 mL Eppendorf. After 4 min of settling, the bottom 50 μL of the biomass was selected for and placed in fresh media for the next growth phase. To increase the oxygen concentration in the supplementary oxygen treatment, air was blown into the tubes directly in the cultures^12^. We measured the oxygen present in the cultures over time using a fiber-optic oxygen optode (FireStingO2, PyroScience, GmbH, Germany).

### Measuring cluster and population size

Cell, cluster and population size distributions were calculated from images of populations of clusters taken in 24-well plates on Nikon Eclipse Ti-E microscope, with composite images taken from a 5×7 tiling array at 40x magnification. To measure the size of multicellular groups, we generated an automated image segmentation pipeline using ImageJ and Matlab, which allowed us to measure the cross-sectional area of individual clusters.

### Monoculture evolutionary replay experiment to examine character displacement

To determine if the difference in size between large and small genotypes is due to character displacement, we evolved four replicate populations of the Large and Small isolate in monoculture for 40 days, with identical media and settling selection conditions as the original MuLTEE experiment^12^. We imaged an average 1224 clusters of each population on days 0, 5, 10, 15, 20 and 40. Information about cluster radius size was obtained by running the images through a Matlab pipeline.

### Sequencing data analysis

In order to quantify the genetic divergence between the Small and Large phenotypes, we sequenced three Large and three Small isolates from lime PO-4 after 715 days of evolution as well as the ancestor. We extracted the DNA using the Life Science Yeast Genomic DNA Purification Kit (VWR). We sent the samples for library preparation and sequencing at the Microbial Genome Sequencing Center (MiGS, https://www.migscenter.com). Illumina Nextera kit (Illumina, San Diego, CA) was used for library preparation and the samples were sequenced with NextSeq 550 platform. We checked the 150 base pairs paired end reads quality using FastQC (version 0.11.5, https://www.bioinformatics.babraham.ac.uk/projects/fastqc/), and decided to trim the first 15 cycles and the last cycle of the Illumina sequence run using FASTP (version 0.21.0)^62^. We kept reads with a PHRED score <30 with a tolerance of 5% of unqualified base pairs. We aligned the reads on the most recent version of S288c strain reference genome (http://sgd-archive.yeastgenome.org/sequence/S288C_reference/genome_releases/S288C_reference_genome_Current_Release.tgz) using BWA-MEM (version 0.7.17-r1188)^63^. We used PICARD (version 2.24.0, http://broadinstitute.github.io/picard/), SAMTOOLS (version 1.7)^64^, BAMTOOLS (version 2.3.0)^65^ to sort, index, remove duplicates and convert to bam the aligned reads. We removed the duplicated reads before performing a quality check of the bam files with GATK ValidateSamFiles tool (GATK version 4.1.9.0). GATK HaplotypeCaller was used on the fixed bam files to call variants^66^. Next, we filtered out the low quality variants by applying the following criteria on VCF files: minimum allele frequency of 0.1, quality score of 30, minimum depth of 12 and maximum depth of 350 using VCFtools (0.1.17)^67^. We compared the variants found in the ancestor to the variants found in the evolved lines to filter out the variants that were already present in the ancestor. All mutations were visually checked on Integrative Genomics Viewer (IGV, version 2.8.13)^68^ and filtered based on the position (telomeric or not), the quality of alignment files of both the ancestor and the evolved samples. We used Beast2 (Bayesian Evolutionary Analysis Sampling Trees) to build the phylogeny with the Jukes-Cantor substitution model and a strict clock^69, 70^.

### Model formulation

#### a) Oxygen diffusion within the clusters

Throughout the culture cycle oxygen is the main limiting factor for the growth of yeast cells. Due to their obligate aerobic metabolism, cells in the cluster that do not have access to oxygen cannot grow. The depth at which oxygen can travel and thus the proportion of cells within the cluster that can grow is dependent on cluster size, the cluster packing fraction, and the resource concentration. We model the oxygen concentration *R*(*r*, *t*) across the diameter of a yeast cluster. The dynamics are dictated by the diffusion, in spherical coordinates, from outside of the cluster of radius *L* by the boundary condition *R*(*L*, *t*) = *R*_0_, where *R*_0_ is the equilibrium concentration of oxygen in the media. Oxygen is then depleted linearly at a rate *γ*, scaled by the local yeast packing fraction *φ*. The partial differential equation that governs this behavior is then:

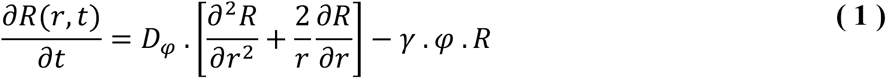

The effective diffusion coefficient *D_φ_* is given by:

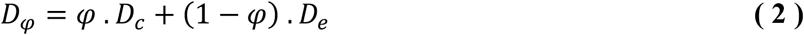

Where the different diffusion coefficients for the yeast cluster *D*_φ_= 17.1 μm^2^s^−1 71^ and the environment *D_e_* = 2000 μm^2^s^−1 72^, are related through the packing fraction *φ*. We used previously published data^12, 17^ to estimate the parameter values for *γ* and the maximum packing fraction *φ*.

One important control parameter for oxygen uptake is how yeast cells are distributed throughout the cluster. We measured the distribution of yeast cells in 20 different clusters using scanning electron microscopy. We used a Zeiss Sigma VP 3View scanning electron microscope (SEM) equipped with a Gatan 3View SBF microtome installed inside a Gemini SEM column to obtain high resolution images of the internal structure of snowflake yeast groups and locate the positions of all cells. All SEM images were obtained in collaboration with the University of Illinois’s Materials Research Laboratory at the Grainger College of Engineering. Snowflake yeast clusters were grown overnight in YPD media, then fixed, stained with osmium tetroxide, and embedded in resin in an Eppendorf tube. A cube of resin 200μm x 200μm x 200μm was cut out of the resin block for imaging. The top surface of the cube was scanned by the SEM to acquire an image with resolution 50nm per pixel (4000 × 4000 pixels). Then, a microtome shaved a 50-nm-thick layer from the top of the specimen, and the new top surface was scanned. This process was repeated until 4000 images were obtained so that the data cube had equal resolution in x,y,z dimensions. Custom image analysis scripts in MATLAB segmented and located the cells. From this data, we calculated the local packing fraction through Voronoi tessellation, finding that the maximum local packing fraction was about 0.6^73^, which corroborated our estimation (see above). Next, for each cluster, we measured the number of cells encountered as a function of distance from the edge of the cluster (Figure 3B). We modeled this distribution as a normal distribution with scale parameter *σ* from the average radius (17μm) and the standard deviation (3μm) as follows: 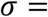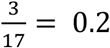. Making the simplifying assumption that the cell distribution within clusters is the same regardless of the cluster size, we used these parameters to calculate the diffusion depth for oxygen for a range of oxygen concentrations and cluster sizes following equations (1) and (2). See section d) for a table of all parameter values.

#### b) Population and resource dynamics

Population dynamics for Small (*S*) and Large (*L*), defined in terms of the number of groups of each genotype, are described using a two-resource Monod growth equation^25^, as follows:

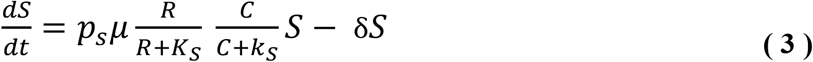

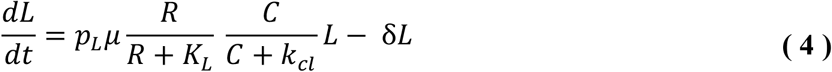

where *R* is the % of oxygen, *C* is the % of carbon, *K* and *k* are the half saturation constants for oxygen and carbon, respectively. *μ* is the maximum cluster growth rate, *P_S_* and *P_L_* are the proportion of active cells within each Small and Large cluster that we calculated using the partial differential equations (section a)), and δ is the basal mortality rate of cells within each cluster. We used a published maximum possible growth rate, *μ*, as the rate of unicellular yeast division in YPG, equal to 0.4 h^−1 74^. We set the mortality rate δ, estimated from previous measurements in snowflake yeast^13^, to 0.008 per hour, representing ~2% per day. The concentration of oxygen dissolved in the media, relative to 100% saturation at present atmospheric levels (PAL), is denoted *R* and is the most critical resource limiting yeast growth. It is added in the system and consumed as follows:

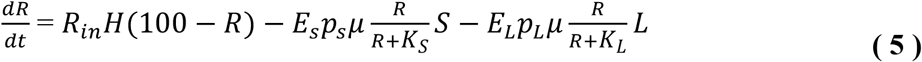

where the addition of new oxygen occurred continuously and is modeled as a Heaviside function:

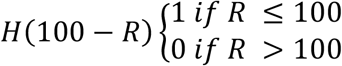

proportional to a % input rate *Rin* (*Rin* = 100, represents the amount of oxygen required to fully oxygenate an entirely anoxic population). Finally, yeast also consume glycerol, which eventually becomes the limiting resource when oxygen is supplemented near the end of the 24 h culture phase. We added an equation describing consumption of this carbon source (*C*):

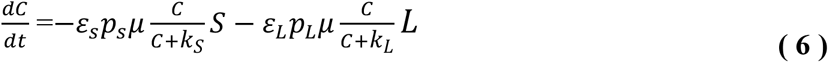

Snowflake yeast convert oxygen and sugar to growth at a rate proportional to their group size, as a larger cluster will require more resource to double than a smaller cluster. This is reflected in the resource utilization variables (*E* and *ε*, for oxygen and carbon), which values linearly increase as a function of size.

#### c) Selection phase due to gravity

In our model, snowflake yeast grow for 24 hours, then the clusters are subject to settling selection where we keep a small proportion (~prior work^17^ has shown that ~1/27^th^ of the population transferred from the bottom of the tube after settling selection to the next tube of fresh media) to initiate the next growth phase. We use Stokes’ law of velocity to estimate the settling selection speed of the clusters as follows:

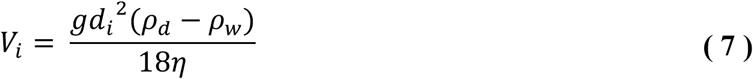

where *d_i_*, is the diameter of Small or Large clusters, *g* is the gravity constant (*g*=9.81 m s^−2^), *ρ_d_* is the mass density of the snowflake yeast (yeast *ρ_d_* =1112.6 kg m^3^), *ρ_w_* is the mass density of water at 25°C,(*ρ_w_* =997 kg m^3^) and *η* is the dynamic viscosity of water at 25°C (*η* =9.81 x 10^−4^ Pa. s^−1^)^26^.

#### d) Model parameters meaning, values and units

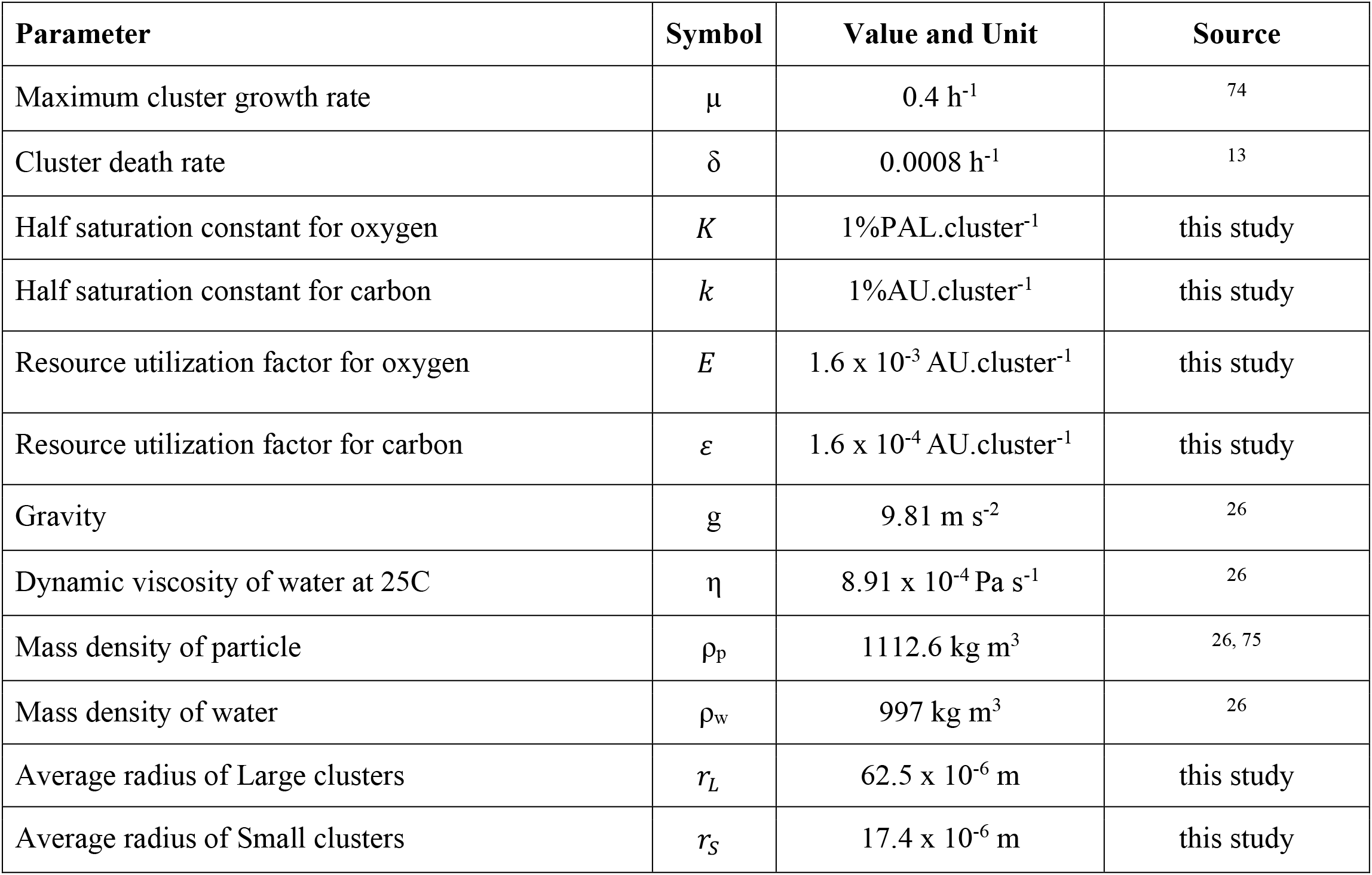
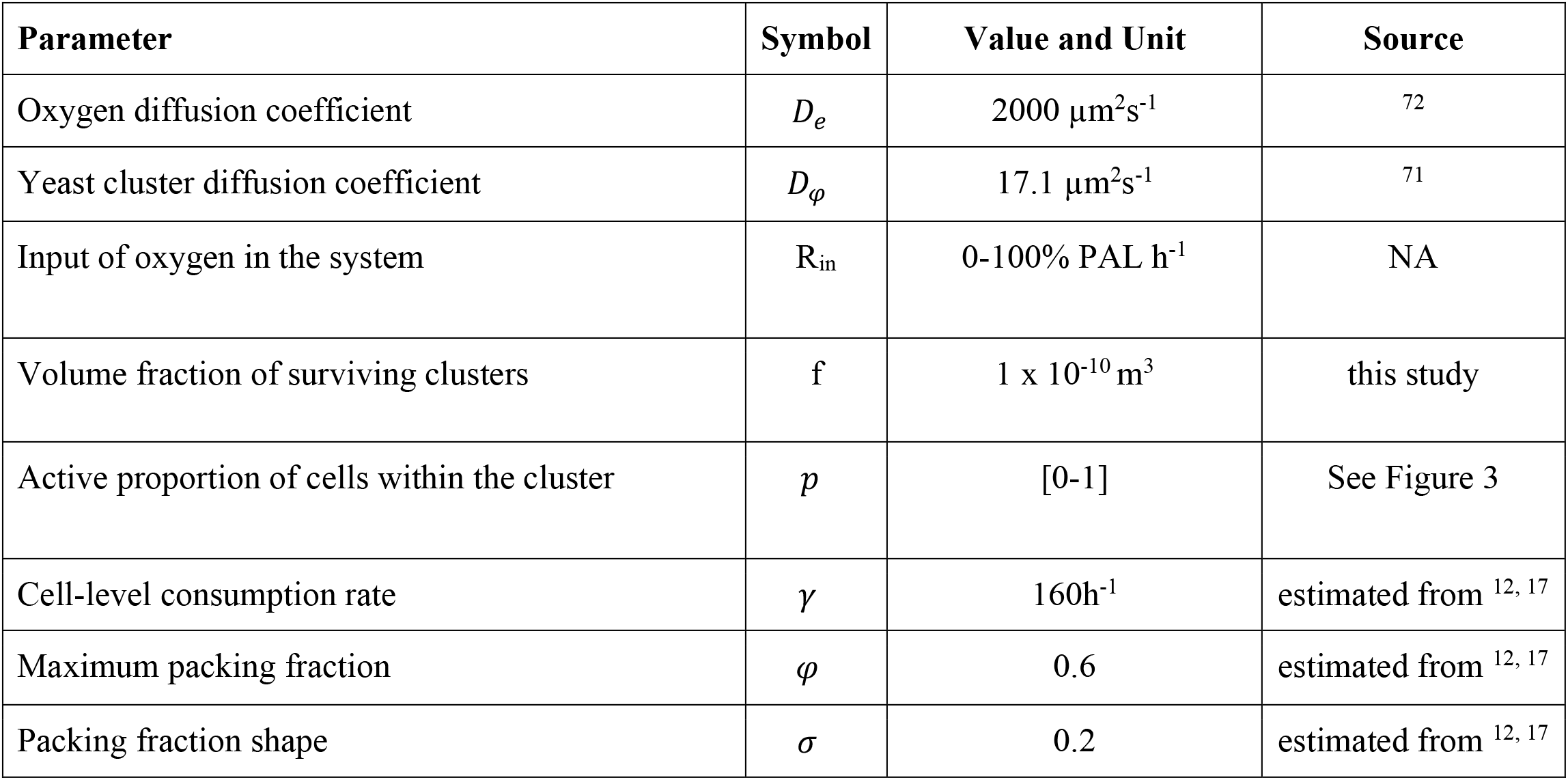

### Simulations of size evolution and coexistence

To examine the state space under which diverse sized snowflake yeast coexist, or when only one size wins the competition (Figure 5), we performed a competitive sweep and looked for evidence of frequency-dependent selection driving coexistence. We performed these competitions for genotypes with sizes ranging from 10-100 μm, in 1 μm increments (828 comparisons total). We started these competitions at increasing initial frequencies for each competing pair, from 1% to 99% (1% increment, 99 frequencies tested per strain pair). After the competition reached a stable equilibrium, or one strain drove the other extinct, we recorded the frequency change of the resident population between the start and the end of the simulation for every starting frequency. If one strain always won at all frequencies, then we simply plotted it as Large or Small winning in Figure 5. Alternatively, we marked the outcome as coexistence in Figure 5 if there were positive frequencies of both Small and Large at the end of the simulation.

The pairwise comparisons performed above provide broad overview of how selection acts on size. To see how this might affect the evolution of a lineage across sequential rounds of mutation and selection, we performed an additional simulation tracking initially monomorphic populations that started out either smaller (10 μm) or larger (90 μm) than the size range in which we saw coexistence. We allowed the starting strain to come into size equilibrium given growth and settling selection, then introduced a competitor at a 1% frequency via mutation—it was chosen from a random uniform distribution ± 12 μm in size from the resident strain’s size (this step size was chosen to be large enough to allow the progression of individual mutations to be clear on the plot). We then competed these strains, as above. If one strain outcompeted the other, we iterated the process of mutation and selection until two strains coexisted stably. As seen in Figure 5, this resulted in a rapid convergence on intermediate sized strains capable of coexistence.

## Acknowledgements

We would like to thank the members of the Ratcliff lab for their input on the project, as well as the GT QBioS Graduate Program for the support. This work was supported by grants from the NIH (Grant No. 5R35GM138030) and the NSF Division of Environmental Biology (Grant No. DEB-1845363) to WCR.

## Author contributions

RMP, WCR and GOB designed the study and wrote the paper. RMP and GOB performed the experiments and analyzed the data. TCD performed the SEM experiment. RMP, DD, WCR, EL and JSW developed and analyzed the ODE model. PB and PJY developed and analyzed the PDE model. All authors contributed to editing the paper.

## References

1. McMahon, S. & Parnell, J. The deep history of Earth’s biomass. Journal of the Geological Society 175, 716–720 (2018).

2. Maloof, A.C. et al. The earliest Cambrian record of animals and ocean geochemical change. GSA Bulletin 122, 1731–1774 (2010).

3. Delwiche, Charles F. & Cooper, Endymion D. The Evolutionary Origin of a Terrestrial Flora. Current Biology 25, R899-R910 (2015).

4. Odling-Smee, F.J., Laland, K.N. & Feldman, M.W. Niche Construction. The American Naturalist 147, 641–648 (1996).

5. Kenrick, P. & Strullu-Derrien, C. The origin and early evolution of roots. Plant physiology 166, 570–580 (2014).

6. Boyce, C.K. The evolutionary history of roots and leaves, in Vascular transport in plants 479-499 (Elsevier, 2005).

7. Lucas, W.J. et al. The plant vascular system: evolution, development and functions f. Journal of integrative plant biology 55, 294–388 (2013).

8. Rose, C. & Hammerschmidt, K. What do we mean by multicellularity? The Evolutionary Transitions Framework provides answers. Frontiers in Ecology and Evolution 9(2021).

9. Bonner, J.T. The origins of multicellularity. Integrative Biology: Issues, News, and Reviews: Published in Association with The Society for Integrative and Comparative Biology 1, 27–36 (1998).

10. Bourrat, P., Doulcier, G., Rose, C.J., Rainey, P.B. & Hammerschmidt, K. Tradeoff breaking as model of evolutionary transitions in individuality and the limits of the fitness decoupling metaphor. bioRxiv, 2021.2009.2001.458526 (2022).

11. Ratcliff, W.C., Denison, R.F., Borrello, M. & Travisano, M. Experimental evolution of multicellularity. Proceedings of the National Academy of Sciences 109, 1595–1600 (2012).

12. Bozdag, G.O., Libby, E., Pineau, R., Reinhard, C.T. & Ratcliff, W.C. Oxygen suppression of macroscopic multicellularity. Nature communications 12, 1–10 (2021).

13. Pentz, J.T., Taylor, B.P. & Ratcliff, W.C. Apoptosis in snowflake yeast: novel trait, or side effect of toxic waste? Journal of The Royal Society Interface 13, 20160121 (2016).

14. Jacobeen, S. et al. Cellular packing, mechanical stress and the evolution of multicellularity. Nature physics 14, 286–290 (2018).

15. Dahaj, S.A.Z. et al. Spontaneous emergence of multicellular heritability. bioRxiv, 2021.2007.2019.452990 (2021).

16. Ratcliff, W.C., Fankhauser, J.D., Rogers, D.W., Greig, D. & Travisano, M. Origins of multicellular evolvability in snowflake yeast. Nature communications 6, 1–9 (2015).

17. Bozdag, G.O. et al. De novo evolution of macroscopic multicellularity. bioRxiv (2021).

18. Tong, K., Bozdag, G.O. & Ratcliff, W.C. Selective drivers of simple multicellularity. Current Opinion in Microbiology 67, 102141 (2022).

19. Knoll, A.H. The multiple origins of complex multicellularity. Annual Review of Earth and Planetary Sciences 39, 217–239 (2011).

20. Herron, M.D. et al. De novo origins of multicellularity in response to predation. Scientific Reports 9, 2328 (2019).

21. Kapsetaki, S.E. & West, S.A. The costs and benefits of multicellular group formation in algae*. Evolution 73, 1296–1308 (2019).

22. Rebolleda-Gómez, M., Ratcliff, W.C., Fankhauser, J. & Travisano, M. Evolution of simple multicellularity increases environmental complexity. Biorxiv, 067991 (2016).

23. Levin, B. Frequency-dependent selection in bacterial populations. Philosophical Transactions of the Royal Society of London. B, Biological Sciences 319, 459–472 (1988).

24. Heino, M., Metz, J.A.J. & Kaitala, V. The enigma of frequency-dependent selection. Trends in Ecology & Evolution 13, 367–370 (1998).

25. Monod, J. The growth of bacterial cultures. Annual review of microbiology 3, 371–394 (1949).

26. Ratcliff, W.C., Pentz, J.T. & Travisano, M. Tempo and mode of multicellular adaptation in experimentally evolved Saccharomyces cerevisiae. Evolution 67, 1573–1581 (2013).

27. Schirrmeister, B.E., de Vos, J.M., Antonelli, A. & Bagheri, H.C. Evolution of multicellularity coincided with increased diversification of cyanobacteria and the Great Oxidation Event. Proceedings of the National Academy of Sciences 110, 1791–1796 (2013).

28. Szathmáry, E. & Smith, J.M. The major evolutionary transitions. Nature 374, 227–232 (1995).

29. Szathmáry, E. Toward major evolutionary transitions theory 2.0. Proceedings of the National Academy of Sciences 112, 10104–10111 (2015).

30. Simpson, G.G. Tempo and mode in evolution. (Columbia University Press, 1944).

31. Ratcliff, W.C., Herron, M., Conlin, P.L. & Libby, E. Nascent life cycles and the emergence of higher-level individuality. Philosophical Transactions of the Royal Society B: Biological Sciences 372, 20160420 (2017).

32. Yoder, J. et al. Ecological opportunity and the origin of adaptive radiations. Journal of evolutionary biology 23, 1581–1596 (2010).

33. Le Hir, G. et al. The climate change caused by the land plant invasion in the Devonian. Earth and Planetary Science Letters 310, 203–212 (2011).

34. Dahl, T.W. & Arens, S.K. The impacts of land plant evolution on Earth’s climate and oxygenation state–an interdisciplinary review. Chemical Geology 547, 119665 (2020).

35. Erwin, D.H. & Tweedt, S. Ecological drivers of the Ediacaran-Cambrian diversification of Metazoa. Evolutionary Ecology 26, 417–433 (2012).

36. Warren, L. et al. Origin and impact of the oldest metazoan bioclastic sediments. Geology 41, 507–510 (2013).

37. McIlroy, D. & Logan, G.A. The impact of bioturbation on infaunal ecology and evolution during the Proterozoic-Cambrian transition. Palaios 14, 58–72 (1999).

38. Billingham, J. Life in the universe: proceedings of a conference held at NASA AMES Research Center, Moffett Field, California, June 19-20, 1979, Vol. 2156. (National Aeronautics and Space Administration, Scientific and Technical …, 1981).

39. Kassen, R. The experimental evolution of specialists, generalists, and the maintenance of diversity. Journal of evolutionary biology 15, 173–190 (2002).

40. Doebeli, M., Ispolatov, Y. & Simon, B. Point of view: Towards a mechanistic foundation of evolutionary theory. Elife 6, e23804 (2017).

41. Roff, D.A. Life history evolution, Vol. 7. (Sinauer Associates Sunderland, MA, 2002).

42. Stearns, S.C. Trade-offs in life-history evolution. Functional ecology 3, 259–268 (1989).

43. Sibly, R. (Oxford Univ Press Oxford, 2002).

44. Rainey, P.B. & Travisano, M. Adaptive radiation in a heterogeneous environment. Nature 394, 69–72 (1998).

45. Koza, A., Moshynets, O., Otten, W. & Spiers, A.J. Environmental modification and niche construction: developing O2 gradients drive the evolution of the Wrinkly Spreader. The ISME Journal 5, 665–673 (2011).

46. Kinnersley, M. et al. Ex uno plures: clonal reinforcement drives evolution of a simple microbial community. PLoS genetics 10, e1004430 (2014).

47. Frenkel, E.M. et al. Crowded growth leads to the spontaneous evolution of semistable coexistence in laboratory yeast populations. Proceedings of the National Academy of Sciences 112, 11306–11311 (2015).

48. Mizuuchi, R., Furubayashi, T. & Ichihashi, N. Evolutionary transition from a single RNA replicator to a multiple replicator network. Nature Communications 13, 1460 (2022).

49. Mahler, D.L., Revell, L.J., Glor, R.E. & Losos, J.B. Ecological opportunity and the rate of morphological evolution in the diversification of Greater Antillean anoles. Evolution: International Journal of Organic Evolution 64, 2731–2745 (2010).

50. Wellborn, G.A. & Langerhans, R.B. Ecological opportunity and the adaptive diversification of lineages. Ecology and evolution 5, 176–195 (2015).

51. Dobson, A., Tilman, D. & Holt, R.D. Unsolved problems in ecology. (Princeton University Press, 2020).

52. Godfray, H.C.J. & May, R.M. Open questions: are the dynamics of ecological communities predictable? BMC biology 12, 1–3 (2014).

53. Godfrey-Smith, P., Bouchard, F. & Huneman, P. Darwinian individuals. From groups to individuals: evolution and emerging individuality 16, 17 (2013).

54. Bonner, J.T. Why size matters, in Why Size Matters (Princeton University Press, 2011).

55. Bonner, J.T. The evolution of complexity by means of natural selection. (Princeton University Press, 1988).

56. Velicer, G.J. & Yu, Y.-t.N. Evolution of novel cooperative swarming in the bacterium Myxococcus xanthus. Nature 425, 75–78 (2003).

57. Jeon, K.W. Development of Cellular Dependence on Infective Organisms: Micrurgical Studies in Amoebas. Science 176, 1122–1123 (1972).

58. Velicer, G.J., Kroos, L. & Lenski, R.E. Developmental cheating in the social bacterium Myxococcus xanthus. Nature 404, 598–601 (2000).

59. Hart, S.F.M., Pineda, J.M.B., Chen, C.-C., Green, R. & Shou, W. Disentangling strictly self-serving mutations from win-win mutations in a mutualistic microbial community. eLife 8, e44812 (2019).

60. Harcombe, W. Novel cooperation experimentally evolved between species. Evolution: International Journal of Organic Evolution 64, 2166–2172 (2010).

61. Gietz, R.D. & Schiestl, R.H. Quick and easy yeast transformation using the LiAc/SS carrier DNA/PEG method. Nature Protocols 2, 35–37 (2007).

62. Chen, S., Zhou, Y., Chen, Y. & Gu, J. fastp: an ultra-fast all-in-one FASTQ preprocessor. Bioinformatics 34, i884-i890 (2018).

63. Li, H. Aligning sequence reads, clone sequences and assembly contigs with BWA-MEM. arXiv preprint arXiv:1303.3997 (2013).

64. Li, H. et al. The sequence alignment/map format and SAMtools. Bioinformatics 25, 2078–2079 (2009).

65. Barnett, D.W., Garrison, E.K., Quinlan, A.R., Strömberg, M.P. & Marth, G.T. BamTools: a C++ API and toolkit for analyzing and managing BAM files. Bioinformatics 27, 1691–1692 (2011).

66. Van der Auwera, G.A. et al. From FastQ data to high-confidence variant calls: the genome analysis toolkit best practices pipeline. Current protocols in bioinformatics 43, 11.10. 11-11.10. 33 (2013).

67. Danecek, P. et al. The variant call format and VCFtools. Bioinformatics 27, 2156–2158 (2011).

68. Robinson, J.T. et al. Integrative genomics viewer. Nature biotechnology 29, 24–26 (2011).

69. Jukes, T.H. & Cantor, C.R. CHAPTER 24 - Evolution of Protein Molecules, in Mammalian Protein Metabolism. (ed. H.N. Munro) 21-132 (Academic Press, 1969).

70. Bouckaert, R. et al. BEAST 2: a software platform for Bayesian evolutionary analysis. PLoS computational biology 10, e1003537 (2014).

71. Vicente, A.A., Dluhý, M., Ferreira, E.C., Mota, M. & Teixeira, J.A. Mass transfer properties of glucose and O2 in Saccharomyces cerevisiae flocs. Biochemical Engineering Journal 2, 35–43 (1998).

72. Haynes, W.M., Lide, D.R. & Bruno, T.J. CRC handbook of chemistry and physics. (CRC press, 2016).

73. Day, T.C. et al. Cellular organization in lab-evolved and extant multicellular species obeys a maximum entropy law. Elife 11, e72707 (2022).

74. Tyson, C.B., Lord, P.G. & Wheals, A.E. Dependency of size of Saccharomyces cerevisiae cells on growth rate. Journal of bacteriology 138, 92–98 (1979).

75. Bryan, A.K., Goranov, A., Amon, A. & Manalis, S.R. Measurement of mass, density, and volume during the cell cycle of yeast. Proceedings of the National Academy of Sciences 107, 999–1004 (2010).

