## Supplementary Material for "Emergence and maintenance of stable coexistence during a long-term multicellular evolution experiment"

**Supplementary material, Pineau *et al.***

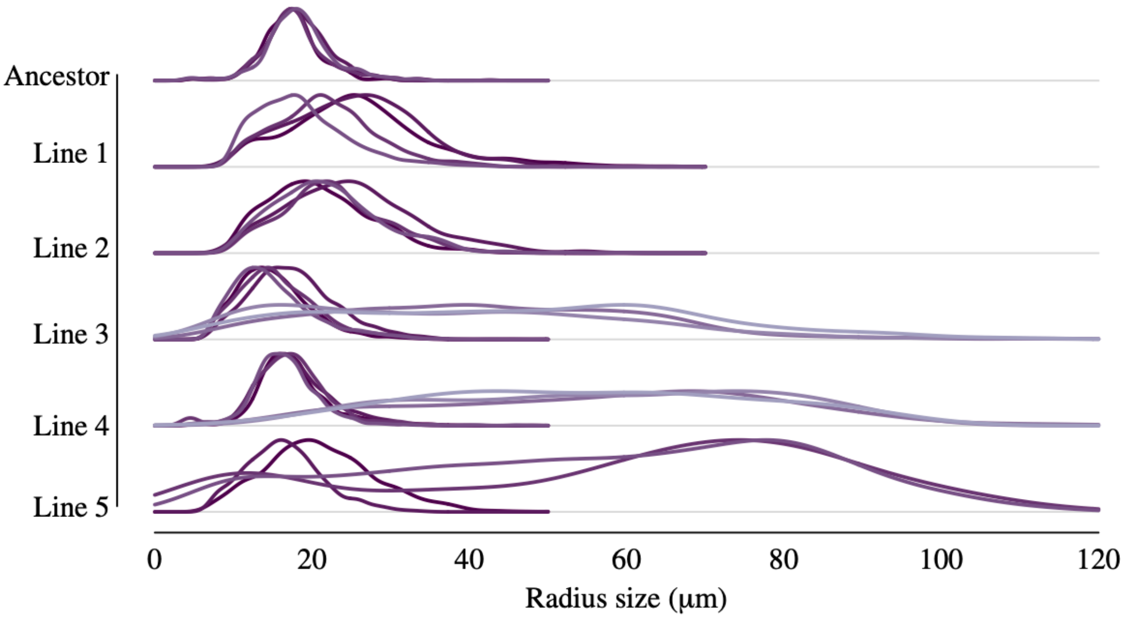


**Supplementary Figure 1**: Size distributions of isolates from the ancestor and the five lines subject to long term evolution, after 715 days of serial transfers (~4,300 generations). We observe the emergence of phenotypic diversity in lines PO-3, PO-4 and PO-5. The different colors denote different isolates.

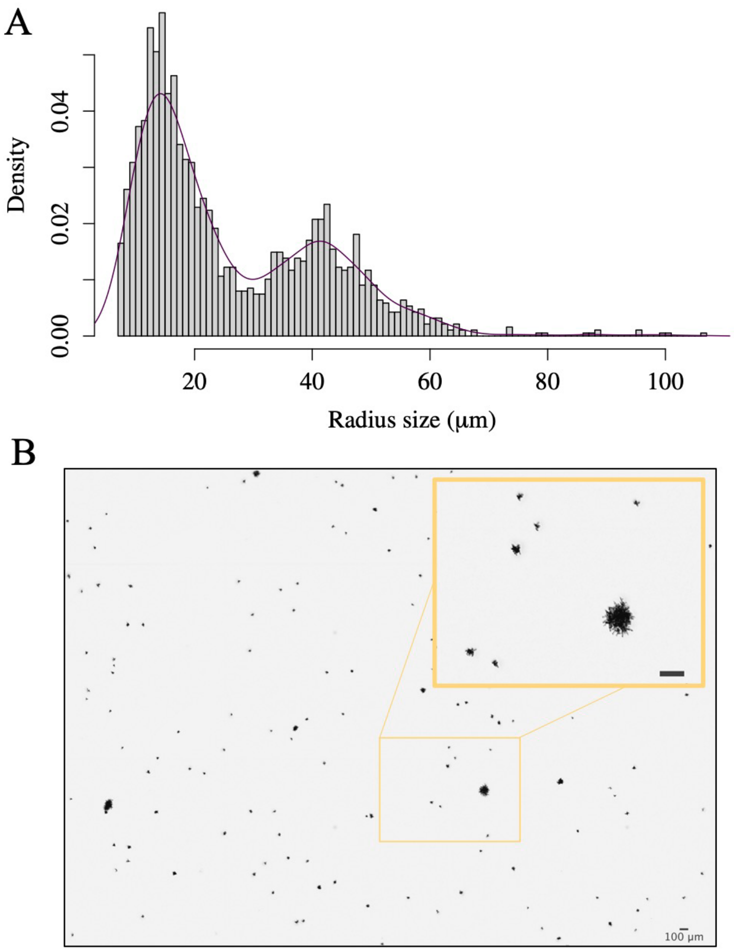


**Supplementary Figure 2**: (A) Whole population size distribution and (B) picture of PO-4 after 715 days of serial transfers (~4300 generations). Large-sized snowflake yeast were present at a mean frequency of 9.4% ± 2.2 in the whole t715 population.

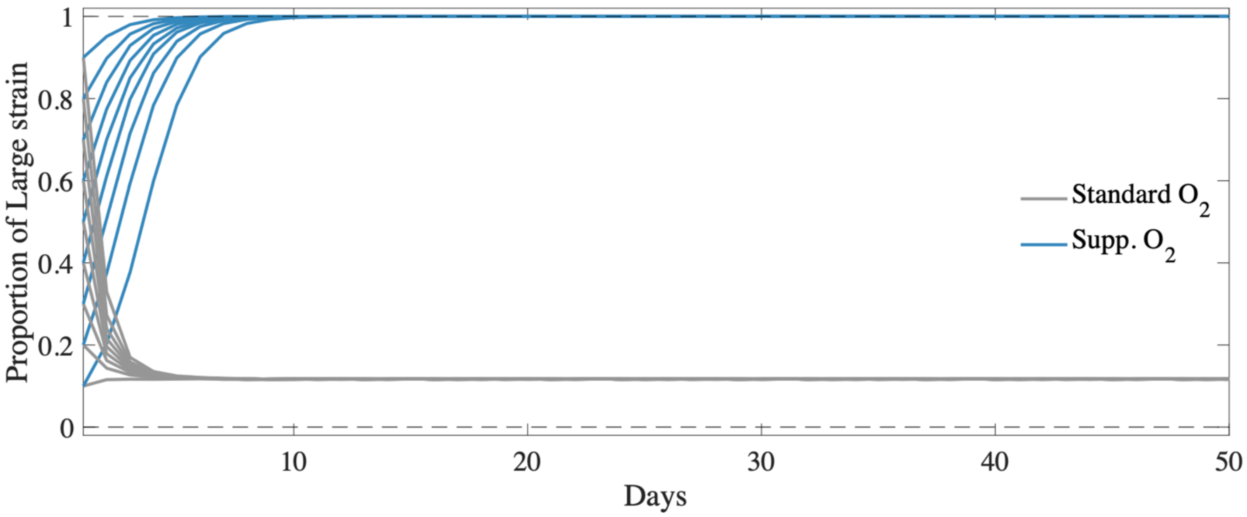


**Supplementary Figure 3**: Long-term dynamics of the competition between Large and Small phenotypes (first 7 days are shown in Figure 2D), in standard (gray) and supplemented oxygen conditions (blue). Coexistence is maintained in standard oxygen conditions. The Large phenotype excludes the Small phenotype when large amounts of oxygen are added into the environment.

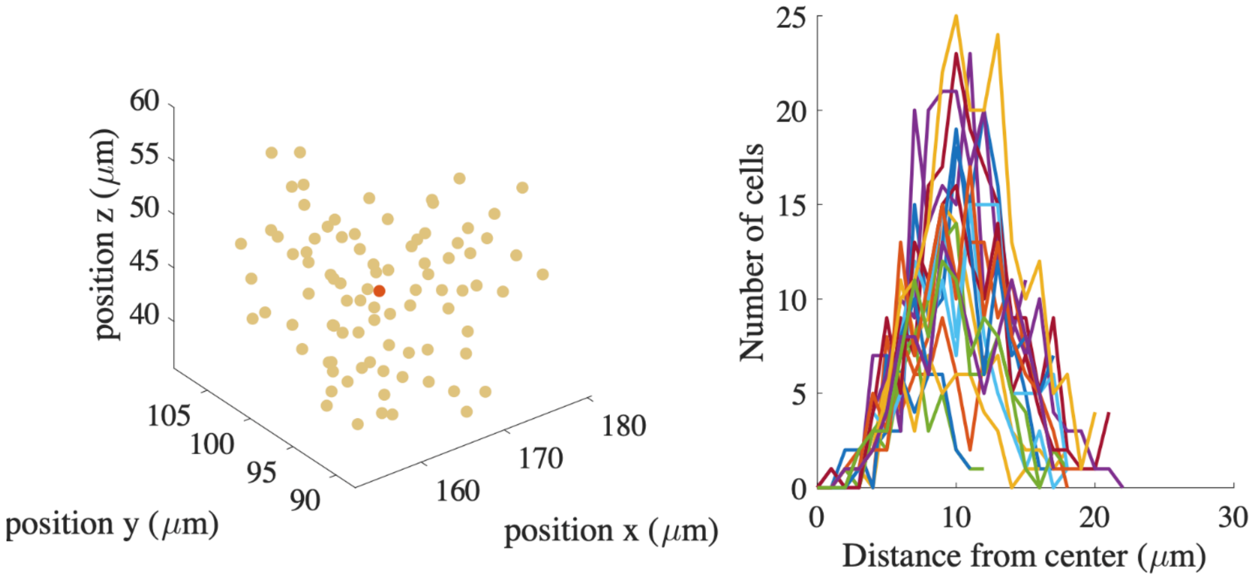


**Supplementary Figure 4**: (left panel) Cell center positions in one example cluster after scanning electron microscopy and image analysis. The red dot shows the center of mass of the cluster. (right panel) Cell number distribution as a function of the distance from the center of the cluster. The cumulative distribution function is shown in Figure 3B in the main text.

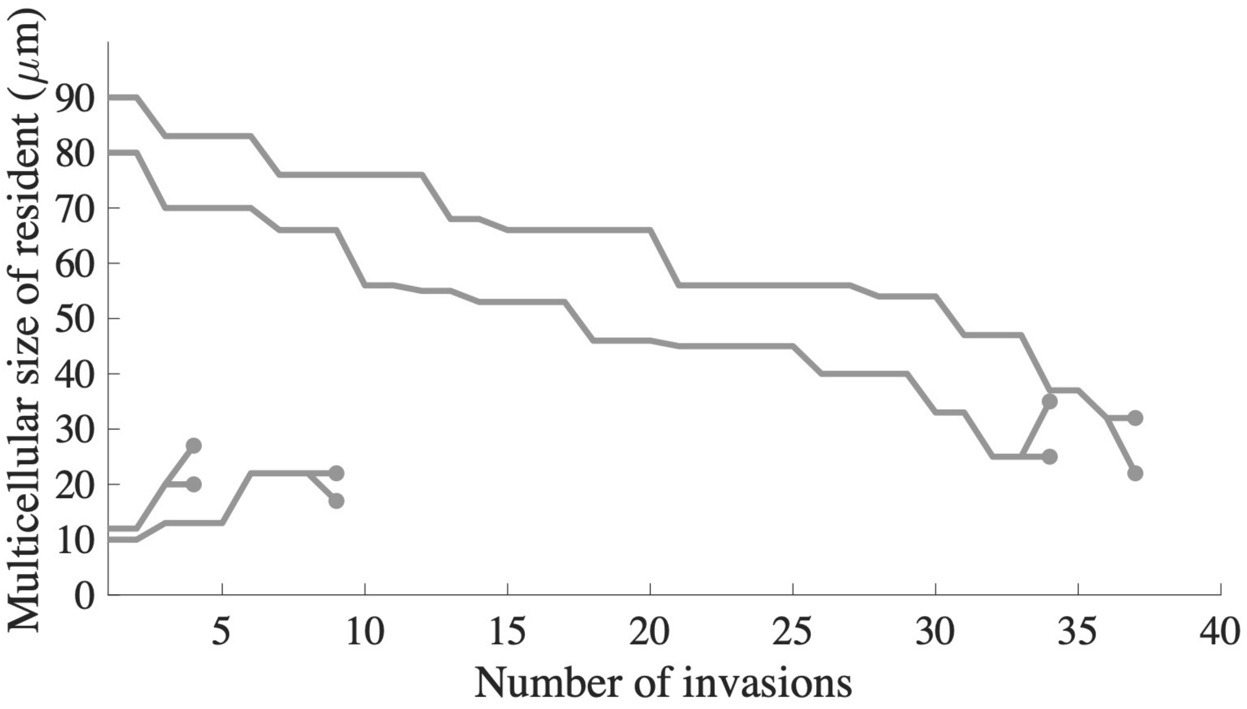


**Supplementary Figure 5**: Example trajectories of the size of resident snowflake yeast when subject to invasion in the model (from Figure 5 simulations). As in the monoculture evolution in Figure 6A, we observe an increase/decrease in size with every invasion until the population reaches the zone of stable polymorphism (dark teal zone in Figure 5), here marked with circles at the end of the trajectories.

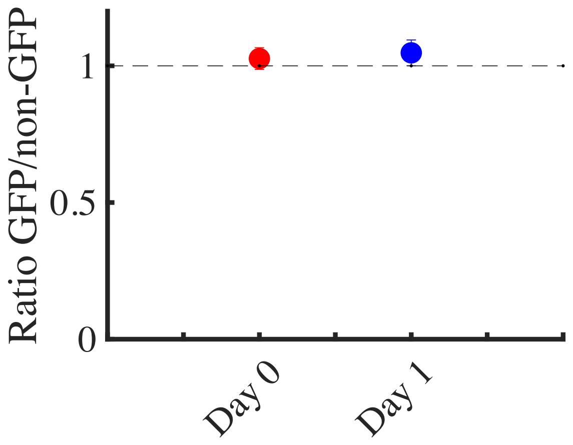


**Supplementary Figure 6**: To measure the fitness cost of GFP expression, we competed a constitutively-expressing GFP unicellular ancestor against a non-GFP control for two days. We did not detect a cost to GFP expression.

**Supplementary Table 1**: Table of the mutations found in 3 Small isolates and 3 Large isolates from the same evolving PO4 line at 715 transfers.
